# High-throughput Screening with Pluripotent Stem Cells Identifies CUDC-907 as an Effective Compound for Restoring the Proinflammatory Phenotype of Nakajo-Nishimura Syndrome

**DOI:** 10.1101/2020.06.01.113894

**Authors:** Naoya Kase, Madoka Terashima, Akira Ohta, Akira Niwa, Fumiko Honda-Ozaki, Yuri Kawasaki, Tatsutoshi Nakahata, Nobuo Kanazawa, Megumu K. Saito

**Author notes:** Author Contribution N.Kase: Conception and design, collection and assembly of data, manuscript writing, data analysis and interpretation; M.K.S.: Conception and design, manuscript writing and final approval of manuscript; M.T.: Conception and design, collection and assembly of data, data analysis and interpretation; A.O.: Conception and design, data analysis and interpretation; A.N., Y.K. F.H-O. T.N.: Conception and design; N.Kanazawa: Provision of the study material and patients.

## Abstract

Nakajo-Nishimura syndrome (NNS) is an autoinflammatory disorder caused by a homozygous mutations in *PSMB8* gene. The administration of systemic corticosteroids is partially effective, but continuous treatment causes severe side effects. We previously established a pluripotent stem cell (PSC)-derived NNS disease model that reproduces several inflammatory phenotypes including the overproduction of monocyte chemoattractant protein-1 (MCP-1) and interferon gamma-induced protein-10 (IP-10). Here we performed high-throughput compound screening (HTS) using this PSC-derived NNS model to find potential therapeutic candidates and identified CUDC-907 as an effective inhibitor of the release of MCP-1 and IP-10. CUDC-907 did not induce cell death within therapeutic concentrations and was also effective on primary patient cells. Further analysis indicated that the inhibitory effect was post-transcriptional. These findings suggest that HTS with PSC-derived disease models is useful for finding drug candidates for autoinflammatory diseases.

**Significance statement:** In this study, we identified a histone deacetylase inhibitor CUDC-907 as a potential effective compound for ameliorating overproduction of inflammatory chemokines in an autoinflammatory disease named Nakajo-Nishimura syndrome. We performed high-throughput screening using pluripotent stem cell-derived monocytic cell lines. Our data prove the validity of screening system as a versatile platform for seeking candidate compounds for the treatment of congenital immunological disorders associated with monocytic lineage cells.

## Introduction

Proteasome-associated autoinflammatory syndromes (PRAAS) are a recently defined group of autoinflammatory disorders caused by the mutations of proteasome subunits or their assembly factors [1]. PRAAS include diseases such as chronic atypical neutrophilic dermatosis with lipodystrophy and elevated temperature (CANDLE) [2], POMP-related auto-inflammation and immune dysregulation disease (PRAID) [3] and Nakajo-Nishimura syndrome (NNS). NNS is caused by a homozygous mutation in *proteasome subunit beta 8* (*PSMB8*)/LMP-7 gene [4, 5]. The onset of NNS begins in early infancy with patients showing a pernio-like rash, then developing periodic high fever, and eventually lipomuscular atrophy and joint contractures mainly in the upper body. The administration of systemic corticosteroids is partially effective against symptoms such as the rash and fever, but not the other symptoms. Further, because the constitutive administration of steroids causes severe side effects, the prognosis of NNS patients relatively remains poor [6-9].

The proteasome is a highly efficient enzyme complex and essential for the degradation of damaged or unnecessary proteins [10]. Such proteins are mainly degraded by a constitutively expressed proteasome to maintain cellular homeostasis or elude cell stress [11]. On the other hand, the immunoproteasome is upregulated upon stimulation by tumor necrosis factor alpha (TNF-α) and/or interferon gamma (IFN-γ), especially in immune cells. The immunoproteasome is required for processing endogenous peptides that are presented on the cell surface as antigens [11-12] and consists of three types of proteases: β1i, β2i and β5i. β5i is encoded by *PSMB8* gene [13].

Almost all NNS patients have the homozygous c.602G > T mutation in *PSMB8* gene, which causes a G201V amino acid substitution in the β5i subunit [4]. It is previously reported that impaired immunoproteasome activity by this mutation causes an excessive accumulation of ubiquitinated and oxidized proteins and an elevated serum concentration of the proinflammatory cytokines and chemokines interleukin-6 (IL-6), monocyte chemoattractant protein-1 (MCP-1) and interferon gamma-induced protein-10 (IP-10) [4]. To investigate the detailed pathophysiology of NNS, we previously established isogenic pairs (wild-type and mutant *PSMB8* G201) of human pluripotent stem cells (PSCs) with or without the NNS-associated *PSMB8* mutation [14]. We then established an *in vitro* disease model by establishing immortalized myeloid cell lines (MLs) from PSC-derived monocytic cells [15]. Although this model shows an excessive accumulation of ubiquitinated proteins and increased secretion of IL-6, MCP-1 and IP-10, the definitive pathogenic mechanism is unknown [14].

High-throughput compound screening (HTS) has been widely adopted as a strategy of drug discovery for intractable diseases including autoimmune disorders [16]. HTS combined with PSC-derived disease models has provided novel drug candidates for several diseases including amyotrophic lateral sclerosis, fibrodysplasia ossificans progressiva and dysferlinopathy [17-19]. Here we established an HTS system using the PSC-derived MLs as model of NNS (NNS-MLs) to discover effective therapeutic candidates for the disease. The HTS system was optimized to identify small molecule compounds inhibiting the production of MCP-1 and IP-10 from NNS-MLs. We thus identified CUDC-907, a histone deacetylase and PI3K-Akt dual inhibitor [20], as the most potent candidate. CUDC-907 seemingly inhibited the translation of MCP-1 and IP-10, thereby decreasing the secretion of these chemokines. These findings suggest that HTS with PSC-derived disease models is an effective approach for identifying therapeutic candidates for PRAAS.

## Material & Methods

### Ethics

This study was approved by the Ethics Committees of Kyoto University (R0091/G0259) and Wakayama Medical University. Written informed consent was obtained from the patients or their guardians in accordance with the Declaration of Helsinki. The use of human embryonic stem cells (ESCs) was approved by the Ministry of Education, Culture, Sports, Science and Technology (MEXT) of Japan.

### Cell Culture

Fibroblasts were collected from two healthy donors (Healthy#1, Healthy#2) and two patients (NNS#1, NNS#2). These fibroblasts were cultured in Dulbecco’s modified eagle medium (08459-64; Nacalai Tesque) in the presence of 10% fetal calf serum (F0926; Sigma-Aldrich) and passaged via dissociation into single cells using 0.25% Trypsin-EDTA (25200-072; Gibco). Fibroblasts were stimulated with 10 ng/mL TNF-α (210-TA; R&D Systems) and 10 ng/mL IFN-γ (285-IF; R&D Systems) for 72 hours to induce the immunoproteasome before all experiments.

Induced pluripotent stem cells (iPSCs) were established from the fibroblasts of NNS#1 [14]. MLs were established from iPSCs bearing mutant homozygous *PSMB8* (MT-iPS-MLs), their isogenic wild-type counterpart (WT-iPS-MLs), and from KhES1 ESCs bearing the same PSMB8 mutation (MT-ES-MLs) [14]. The MLs were cultured in StemPro™-34 serum-free medium (10639-011; Gibco) containing 2 mM L-glutamine (25030-081; Gibco) in the presence of 50 ng/mL M-CSF (216-MC; R&D Systems) and 50 ng/mL GM-CSF (215-GM; R&D Systems). KhES1 ESCs were kindly provided by Dr. Hirofumi Suemori (Kyoto University, Kyoto, Japan).

### HTS system

We used MT-iPS-MLs and homogeneous time-resolved fluorescence (HTRF)-based chemokine measurements for the HTS system. The compound library is described in **Table 1**. For the 1st and 2nd screenings, 5×10^3^ MT-iPS-MLs were seeded into each well of 384-well plates. The plates were individually prepared for the detection of MCP-1 or IP-10. The cells were treated with compounds at concentrations of 1 μM (1st screening) and 100 nM (2nd screening). Three hours later, 50 ng/mL lipopolysaccharide (LPS) (tlrl-peklps; InvivoGen) for MCP-1 and 100 ng/mL IFN-γ for IP-10 was added for the stimulation. After the subsequent culture for 21 hours, the supernatants were sampled, and the Human CCL2 (MCP-1) Kit (62HCCL2PEG; Cisbio) or Human CXCL10 (IP-10) Kit (62HCX10PEG; Cisbio) was added to detect MCP-1 or IP-10 (**Fig. 1A**). The chemokine concentration was detected by using POWERSCAN^®^4 (DS Pharma Biomedical). Validation of the hit compounds with MT-ES-MLs was performed using the same procedure as the 2nd screening.

**Table 1.** The source of compounds in the HTS.

|  |  |  |
| --- | --- | --- |
| Enzo-FDA<br>(FDA Approved Drugs) | Fixed | 636 |
| ENZO ICCB<br>(World Wide Drugs) | Fixed | 477 |
| Microsource-US Drugs<br>(FDA Approved Drugs) | Fixed | 1,020 |
| Microsource-International Drugs<br>(World Wide Drugs) | Fixed | 238 |
| Sigma LOPAC | Fixed | 1,280 |
| Kinase Inhibitor<br>Cell-Permeable compounds | CiRA Special Selection | 1,254 |
| Bioactive Compounds | CiRA Special Selection | 916 |
| **Total** |  | **5,821** |

**Figure. 1.**
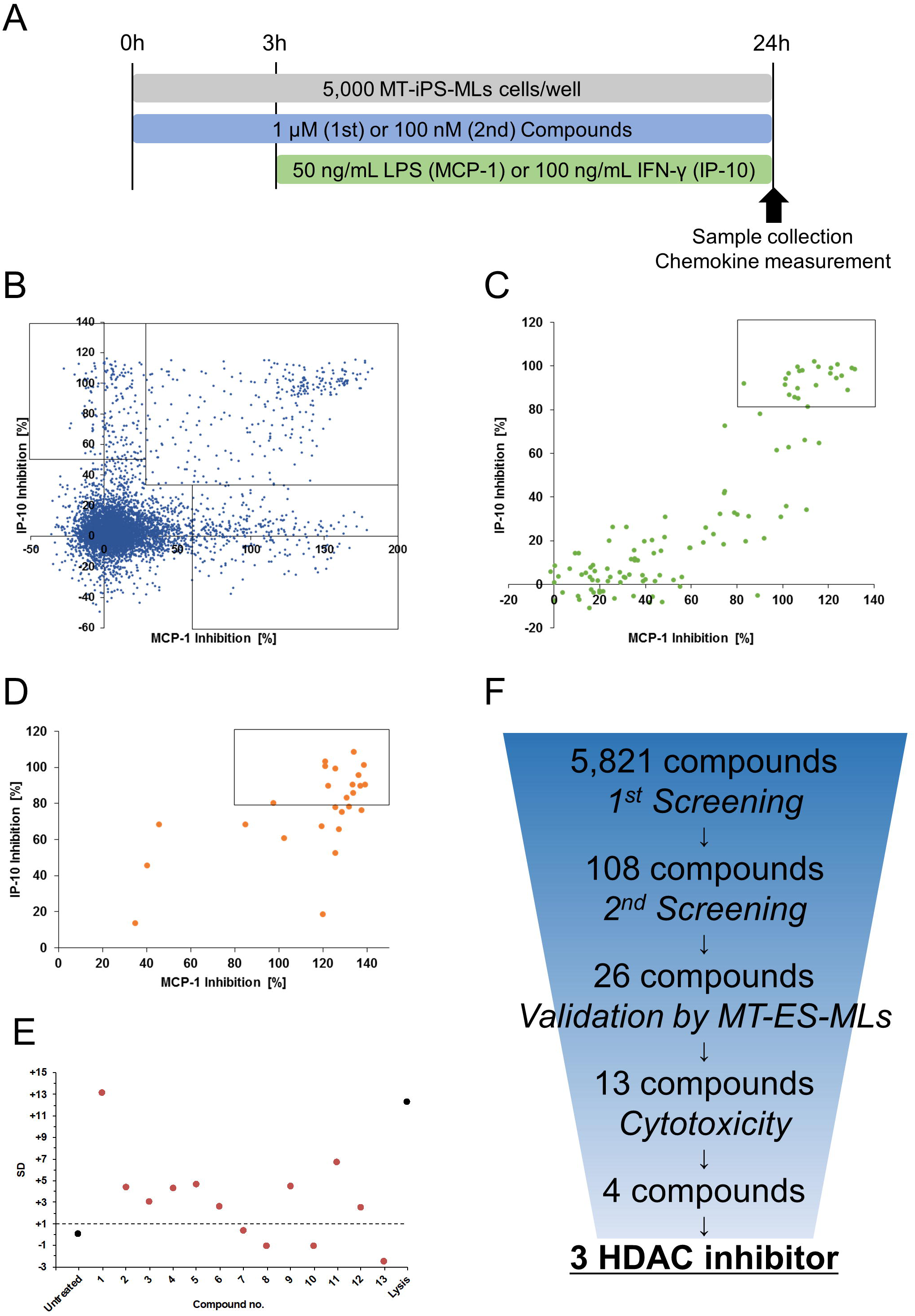
Results of the HTS. (A): Schematic representation of the time course and culture condition for the sampling. (B-C): Results of the 1st (B) and 2nd (C) screening using MT-iPS-MLs. (D) Validation of the hit compounds using MT-ES-MLs. (B-D) Data inside the squares shows hit compounds. (E): Results of the cytotoxic evaluation. Compounds below the dotted line are considered hits. Untreated (DMSO control) and Lysis (100% cell death) are plotted as controls. (F): Schematic representation of the HTS results.

### Enzyme-linked immunosorbent assay (ELISA)

The concentrations of cytokines and chemokines in MLs and fibroblasts were measured by using a LEGENDplex™ Human Adipokine Panel (740196; BioLegend) according to the manufacturer’s protocol. Quantification was done with Guava^®^ easyCyte™ (Luminex). After seeding 1 × 10^4^ MLs or 5 × 10^3^ fibroblasts into each well of 96-well plates, the compounds in DMSO (D2650; Sigma-Aldrich) were applied. Three hours later, TNF-α and IFN-γ at 100 ng/mL for the MLs and 10 ng/mL for the fibroblasts were added for the stimulation. After culturing for 21 hours, the supernatants were sampled.

### Cell cytotoxicity

To evaluate cell cytotoxicity by the extracellular lactate dehydrogenase (LDH) activity, Cytotoxicity Detection Kit^PLUS^ was used according to the manufacturer’s protocol (4744934001; Merck). For MT-ES-MLs, 5×10^3^ cells were seeded into each well of 384-well plates, and then compounds in DMSO at a concentration of 100 nM were applied. Three hours later, 100 ng/mL IFN-γ was added for the stimulation. After culturing for 21 hours, the supernatants were sampled. Detections were performed by using POWERSCAN^®^4. For MT-iPS-MLs and fibroblasts, 1×10^4^ MT-iPS-MLs and 1.5×10^3^ NNS#1 fibroblasts were seeded into each well of 96-well plates. After culturing for 24 hours, the supernatants were sampled. Detections were performed by using 2104 EnVision Multilabel Plate Readers (PerkinElmer). In every case, total cell lysates (100% cell death) from the same number of cells were used as the positive control.

To evaluate cell viability by the intracellular nicotinamide adenine dinucleotide (NADH) activity, 5×10^4^ MT-iPS-MLs or 5×10^3^ NNS#1 fibroblasts were seeded into each well of 96-well plates. After culturing for 24 hours, Cell Counting Kit-8 (CK04; Dojindo) was used according to the manufacturer’s protocol.

### RNA isolation and quantitative polymerase chain reaction (qPCR)

Total RNA was column-purified with the RNeasy^®^ Mini Kit (74106; Qiagen) and treated with RNase-free DNase (79254; Qiagen). Purified RNA was reverse transcribed using PrimeScript™ RT Master Mix (RR037A; Takara) according to the manufacturer’s protocol. qPCR was performed using StepOnePlus™ (Applied Biosystems) with TB Green Premix Ex Taq II (RR820A; Takara). The primer sequences used in this study are listed in **Table 2**.

**Table 2.**
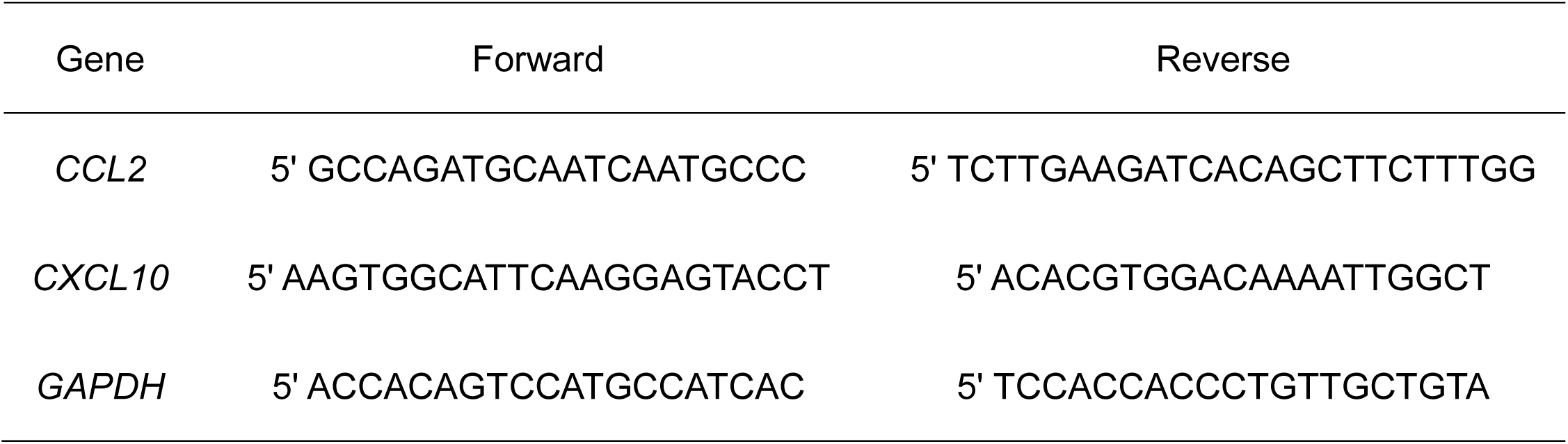
The list of primers for the qPCR.

### Immunoblotting

A total of 3×10^5^ cells were lysed for 30 minutes on ice in RIPA buffer (188-02453; Wako) with protease inhibitor cocktail (04080-11; Nacalai Tesque). After being centrifuged at 15,000g for 5 minutes at 4 °C, the supernatants were collected. The total amount of proteins in lysates was quantified by the *DC*™ Protein Assay (500-0116JA; Bio-Rad) and 2104 EnVision Multilabel Plate Readers (PerkinElmer) according to the manufacturer’s protocol, then the lysates were adjusted to the same concentrations. The lysates were boiled for 5 minutes in 4x Laemmli Sample Buffer (161-0747; Bio-Rad) containing 2-Mercaptoethanol (21418-42; Nacalai Tesque). The proteins were then separated by sodium dodecyl sulfate-polyacrylamide gel electrophoresis and transferred onto an Immobilon-P membrane (IPVH00010; Merck). The membranes were blocked with 10% nonfat milk (190-12865; Wako) in Tris-buffered saline plus 0.1% Tween20 (9005-64-5; Sigma-Aldrich) and reacted with the indicated antibodies. The antibodies used for immunoblotting were as follows: monoclonal anti-GAPDH antibody (2118, CST), polyclonal anti-MCP-1 antibody (ab9669, Abcam), polyclonal anti-IP-10 antibody (ab8098, Abcam), HRP-conjugated anti-rabbit IgG antibody (7074, CST) and HRP-conjugated anti-mouse IgG antibody (7076, CST). SuperSignal™ West Femto Maximum Sensitivity Substrate (34095; Thermo Fisher) was used for the chemiluminescence. Images were obtained using ImageQuant LAS 4000 (GE Healthcare).

### Statistics

All statistical analyses and IC_50_ values were performed using GraphPad Prism (GraphPad Software). Statistical methods are described in the figure legends.

## Results

### HTS identifies HDAC inhibiters as therapeutic candidates for NNS

We performed HTS to identify drug candidates for NNS. Compounds were evaluated by the inhibition rates of MCP-1 and IP-10, both of which are proinflammatory chemokines specifically elevated in NNS patients [5, 14]. Although IL-6 is also specifically elevated in NNS patients, blockage of the IL-6 receptor by the administration of the anti-IL-6 receptor tocilizumab provided limited therapeutic effects [21]. Therefore, we focused on suppressing both MCP-1 and IP-10. The compounds library consisted of 5,821 compounds including approved drugs, kinase inhibitors and bioactive chemicals (**Table 1**). The cells were pretreated with compounds for 3 hours and treated subsequently with LPS or IFN-γ to induce MCP-1 or IP-10, respectively **(Fig. 1A)**. In the 1st and 2nd screenings, we used MT-iPS-MLs. We first evaluated the effect of the 5,821 compounds on MT-iPS-MLs at a concentration of 1 μM. We defined the criteria for hit compounds as >60% inhibition of MCP-1, >50% inhibition of IP-10, or >30% inhibition of both **(Fig. 1B)**. We excluded several hit compounds from the results of the 1st screening because of their potential cytotoxicity. In the 2nd screening, we evaluated 108 compounds at a concentration of 100 nM and picked up those with >80% inhibition of both MCP-1 and IP-10. The 2nd screening yielded 26 hit compounds **(Fig. 1C)**. To validate the reproducibility of the results, we evaluated the inhibitory effects of the 26 compounds on another clone, MT-ES-MLs, at the same 100 nM concentration. We thus obtained 13 hit compounds which inhibited the secretion of both MCP-1 and IP-10 at more than 80% **(Fig. 1D)**. Finally, we evaluated the cytotoxicity of these 13 compounds on MT-ES-MLs at the same concentration. Four compounds showed an LDH release less than one standard deviation (SD) from the mean of the untreated control and were considered non-toxic **(Fig. 1E)**. Since 3 of the 4 compounds are HDAC inhibitors, we conducted detailed comparative studies on these three **(Fig. 1F)**.

### CUDC-907 inhibits MCP-1 and IP-10 production at a lower concentration

We next compared the inhibitory effects of the 3 HDAC inhibitors: CUDC-907 **(Fig. 2A)** [20], JNJ-26481585 **(Fig. 2B)** [22] and LAQ 824 **(Fig. 2C)** [23]. The 50% inhibitory concentration (IC_50_) values were calculated from the dose-dependent effects on MT-iPS-MLs **(Fig. 2D-2I)**. Among the 3 compounds, CUDC-907 showed the lowest IC_50_ for both MCP-1 and IP-10 secretion **(Table 3)**. We therefore focused on CUDC-907 as the first choice for NNS. CUDC-907 is known to be a dual inhibitor of HDAC and PI3K-Akt and is an effective therapeutic candidate for AML, in which the PI3K-Akt pathway is constitutively activated [24]. However, there are no reports yet on its effect on the production of proinflammatory cytokines or chemokines. Since HDAC inhibitors are widely known to arrest the cell cycle [25], we measured the effect of CUDC-907 on cell cytotoxicity by detecting intracellular NADH and extracellular LDH activities. Although CUDC-907 treatment slightly decreased the viability of MT-iPS-MLs in a dose-dependent manner, almost no cytotoxicity was observed within effective concentrations **(Fig. 2J-2K)**. These results suggest that CUDC-907 avoids serious cytotoxic side effects within the effective window and that its inhibitory effect is not attributed to the consequence of cell death.

**Table 3.** IC_50_ values of three HDAC inhibitors. IC_50_ values were calculated from the dose-response curves shown in **Fig. 2D-2I**.

| Compound | MCP-1 (nM) | IP-10 (nM) |
| --- | --- | --- |
| CUDC-907 | 6.0 | 21.7 |
| JNJ-26481585 | 25.7 | 73.6 |
| LAQ824 | 38.6 | 101.9 |

**Figure. 2.**
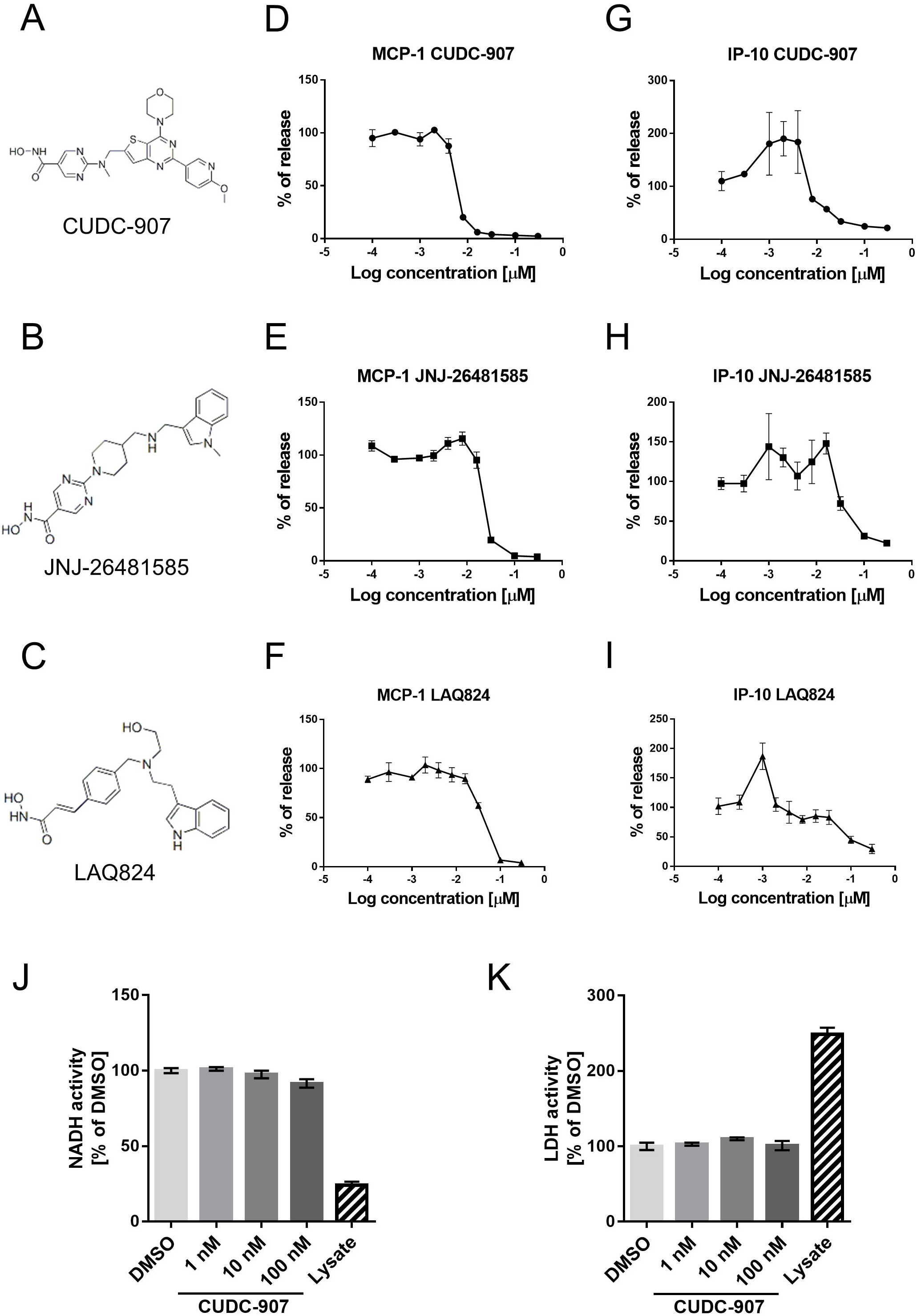
Comparison of three HDAC inhibitors. (A)-(C): Structural formula of each compound. (D)-(I): Dose-dependent inhibitory effects of the compounds on MCP-1 (D)-(F) and IP-10 (G)-(I) (n=3). (J)-(K): Cell cytotoxicity by detecting intracellular NADH (J) and extracellular LDH activities (K) in the presence of CUDC-907 under the indicated concentrations (n=3).

### CUDC-907 exerts effective inhibition in patient fibroblasts

To address the consistency of the inhibitory effects of CUDC-907 on other cell types, we next used primary patient fibroblasts. The expression of the immunoproteasome in fibroblasts is lower than that in immune cells [26]. We therefore measured the production of cytokines after longer treatment (72 hours) with TNF-α and IFN-γ to induce the immunoproteasome sufficiently to show the phenotype of NNS **(Fig. 3A)**. Since the amount of secreted IP-10 was very small in fibroblasts, we could not compare fibroblasts from patients and healthy donors (data not shown). However, the secretion of MCP-1 from patient fibroblasts (NNS#1, NNS#2) was significantly higher than that from healthy donor-derived fibroblasts (Healthy#1, Healthy#2) **(Fig. 3B)**. The IC_50_ of CUDC-907 on the inhibition of MCP-1 secretion was comparable to that in MT-iPS-MLs **(Fig. 3C)**. Furthermore, we measured the cytotoxicity on patient fibroblasts. Almost no cytotoxicity was observed within the effective concentrations **(Fig. 3D-3E)**. Therefore, CUDC-907 inhibits the secretion of MCP-1 in patient fibroblasts at similar concentrations as in MT-iPS-MLs without cell death.

**Figure. 3.**
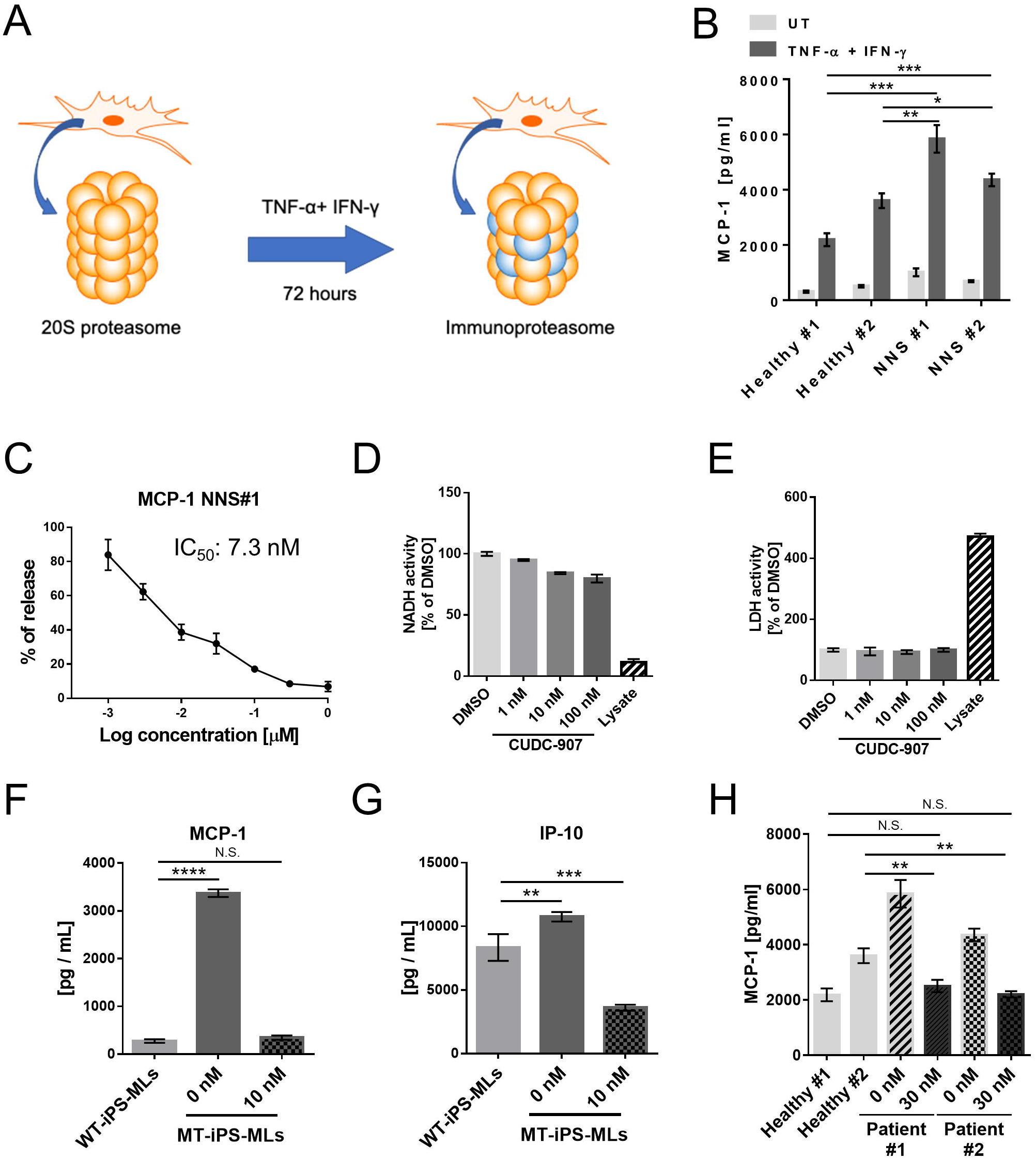
Effects of CUDC-907 on primary patient fibroblasts. (A): Schematic representation of the immunoproteasome induction in fibroblasts. (B): MCP-1 secreted from fibroblasts under TNF-α and IFN-γ treatment (n=3). Statistical analysis was performed by Student’s t test: *p < 0.05; **p < 0.01; ***p< 0.005. (C): Dose-dependent inhibitory effects of CDDC-907 on MCP-1 in NNS#1 fibroblasts. (D)-(E): Cell cytotoxicity of NNS#1 fibroblasts by detecting intracellular NADH (D) and extracellular LDH activities (E) in the presence of CUDC-907 at the indicated concentrations (n=3). (F,G): Secreted MCP-1 (F) and IP-10 (G) from WT-iPS-MLs and MT-iPS-MLs in the presence of CUDC-907 (n=3). Statistical analysis was performed by one-way ANOVA with Dunnett’s multiple comparison: **p < 0.01; ***p < 0.005; ****p< 0.001. (H): Secreted MCP-1 from fibroblasts in the presence of CUDC-907 (n=3). Statistical analysis was performed by Student’s t test: **p < 0.01.

Notably, when we compared the chemokine secretion level of MLs with or without *PSMB8* mutation (MT-iPS-MLs and WT-iPS-MLs, respectively), CUDC-907 sufficiently suppressed the amount of secreted MCP-1 and IP-10 by MT-iPS-ML to levels comparable with those secreted by WT-iPS-MLs **(Fig. 3F and 3G)**. CUDC-907 also suppressed MCP-1 secretion from mutant fibroblasts to a level similar level as wild-type fibroblasts (**Fig. 3H**).

### CUDC-907 inhibits the production of MCP-1 and IP-10 post-transcriptionally

To understand the inhibition mechanism of CUDC-907, we quantified the expression levels of *CCL2* gene encoding MCP-1 and *CXCL10* gene encoding IP-10. Contrary to the results of ELISA, CUDC-907 upregulated the expression of these two genes **(Fig. 4A)**, indicating that CUDC-907 exerts its inhibitory effect post-transcriptionally. In line with this assumption, we next observed the protein levels of MCP-1 and IP-10. To detect intracellular MCP-1 and IP-10 levels by western blot, we inhibited the extracellular release of these cytokines by treating the cells with Brefeldin A, an inhibitor of the protein transportation from endoplasmic reticulum to Golgi apparatus [27]. In the presence of Brefeldin A, the intracellular levels of MCP-1 and IP-10 were increased in all conditions observed (Fig. 4B), indicating Brefeldin A inhibited the extracellular release of MCP-1 and IP-10. In the presence of Brefeldin A, CUDC-907 decreased the intracellular protein levels of MCP-1 and IP-10 (Fig. 4B). These results indicate that CUDC-907 suppresses the production of MCP-1 and IP-10 by inhibiting translational processes.

**Figure. 4.**
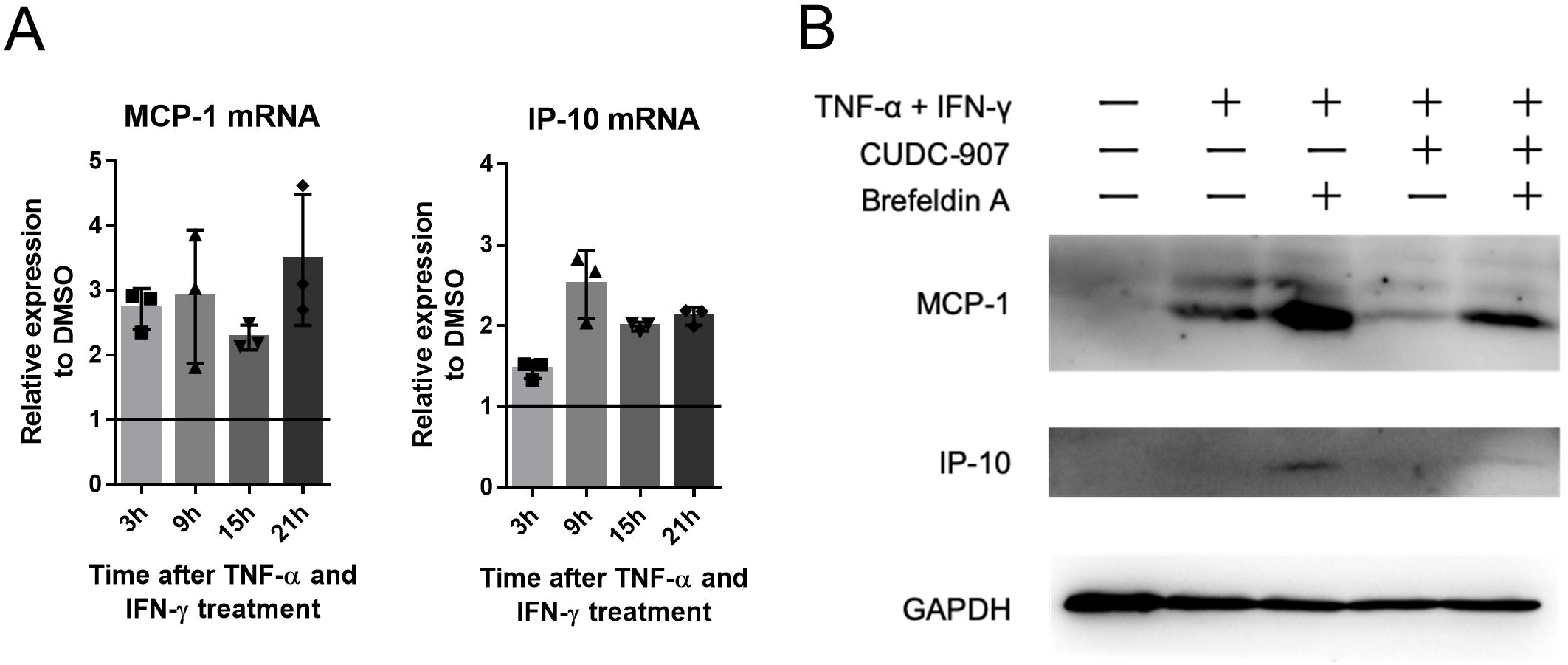
Identification of the inhibitory point of CUDC-907. (A): Relative expression levels of *CCL2* and *CXCL10* genes to DMSO control (n=3). Cells pre-treated with CUDC-907 (10 nM) for 3 h were sampled after treatment with TNF-α and IFN-γ (100 ng/mL) at the indicated times. The expressions were normalized to the *GAPDH* gene expression. (B): Total amount of intracellular MCP-1, IP-10 and glyceraldehyde 3-phosphate dehydrogenase (GAPDH) as the loading control. Cells pre-treated with CUDC-907 (10 nM) for 30 min were sampled after treatment with TNF-α and IFN-γ (100 ng/mL) for 6 h. Brefeldin A was added 1 h before the end of the culture. A representative gel image from three independent experiments is shown.

## Discussion

In this study, we identified an effective compound that restores the disease-associated proinflammatory phenotype of NNS by using PSC-based HTS. We used MLs established from PSCs for the HTS, because MLs are easy to propagate, freeze-stock, and differentiate into functional macrophages or dendritic cells [15]. We previously demonstrated that such MLs can be used to model the proinflammatory phenotypes of various autoinflammatory disorders [14, 28-30]. Combined with the current study, which successfully identified candidate compounds, we concluded that this ML-based HTS system provides a versatile platform for screening candidate compounds for treating autoinflammatory disorders.

MCP-1 and IP-10 are known as proinflammatory chemokines [31-32]. Their increased production is associated with the inflammatory symptoms of various diseases, and their suppression sometimes ameliorates the symptoms in diseases such as polycystic kidney disease and rheumatoid arthritis [33-34]. Therefore, we hypothesized that the suppression of the overproduction could alleviate the autoinflammatory symptoms of NNS. However, NNS patients are known to suffer from lipomuscular atrophy in addition to inflammatory symptoms [6, 9]; whether the suppression of MCP-1 and IP-10 also alleviates these atrophic symptoms is unknown. New lipomuscular models of NNS would help in this regard. PSC-derived muscular disease models have contributed to the elucidation of pathophysiologies and drug discovery [19, 35] and should be of benefit to understanding the pathophysiology of the muscular lesions in NNS. The *in vivo* effects of compounds on animal models should be also evaluated.

CUDC-907 is a dual inhibitor of HDAC and PI3K-Akt [20]. It has also been reported to inhibit the growth of several tumor cells, including multiple myeloma and chronic lymphoma [36-37]. Based on those reports, CUDC-907 has been considered an anticancer drug. However, in this study, we demonstrated for the first time that CUDC-907 has an inhibitory effect on the extracellular release of proinflammatory chemokines. This finding suggests that CUDC-907 could be applied to a variety of inflammatory disorders including NNS. However, given the tumor toxicity of CUDC-907 [36-37], its immediate clinical application to NNS patients is not realistic. Further evaluation of its effects and safety should be performed in the future.

A large number of compounds are reported to inhibit the release of extracellular cytokines or chemokines, many of which suppress the expression levels by targeting the inflammatory signaling cascade [38-39]. However, for some diseases such as asthma and diabetic retinopathy, abnormal post-transcriptional regulation is key [40-41]. For such diseases, the inhibition of the post-transcriptional phase is considered more appropriate in the therapeutic strategy [40-41]. In the present study, CUDC-907 was found to inhibit the production of MCP-1 and IP-10 post-transcriptionally. Future study of the detailed molecular mechanism of this effect could provide therapeutic advances for such diseases.

## Conclusion

In this study, we performed HTS with a PSC-derived NNS disease model to find a potential therapeutic candidate, CUDC-907, which is an HDAC and PI3K-Akt dual inhibitor. CUDC-907 effectively inhibits the release of MCP-1 and IP-10 without inducing cell death and is also effective on primary patient cells. CUDC-907 inhibits the production of MCP-1 and IP-10 post-transcriptionally. These findings suggest that HTS with PSC-derived disease models is an effective approach at identifying therapeutic candidates for autoinflammatory disorders.

## Acknowledgements

We thank Ms. Harumi Watanabe for providing administrative assistance and Dr. Peter Karagiannis for proofreading the paper. Funding was provided from the Core Center for iPS Cell Research of Research Center Network for Realization of Regenerative Medicine from the Japan Agency for Medical Research and Development (AMED) [M.K.S.], the Acceleration Program for Intractable Diseases Research utilizing Disease-specific iPS cells from AMED (17935244 and 15652070) [M.K.S.], Practical Research Project for Rare/Intractable Diseases from AMED (17929899) [M.K.S.], the Translational Research program; Strategic PRomotion for practical application of INnovative medical Technology (TR-SPRINT) from AMED [M.K.S.] and Wakayama Medical University Special Grant-in-Aid for Research Projects [N.K.]

## Disclosure of Potential Conflicts of Interest

The authors declare no conflict of interest.

## Data Availability Statement

The data that support the findings of this study are available from the corresponding authors upon reasonable request.

## Notes

### Competing Interest Statement

The authors have declared no competing interest.

